# Whole genome sequencing and comparative genomic studies of *Priestia filamentosa* JURBA-X for its drought-tolerance, plant-growth promotion, and fluorescent characteristics

**DOI:** 10.1101/2024.04.09.588649

**Authors:** Sneha Murthy, Mallu Govardhana, Kumudini Belur Satyan, Gaurav Sharma

## Abstract

*Priestia filamentosa* JURBA-X is a nonmotile, endospore-forming, and Gram-positive bacterium isolated from a rhizosphere soil sample of groundnut fields in Andhra Pradesh, India, during summer. JURBA-X exhibits chains of filamentous morphology with vibrant yellow fluorescence. It shows tolerance to drought stress, phosphate solubilization, siderophore production, and antibacterial activity. 16s rRNA and single-copy orthologous DNA gyrase subunit B-based phylogenies suggested its closeness with *Priestia filamentosa* spp., leading to its classification as *P. filamentosa* JURBA-X. The whole genome was assembled into 5,113,908 bp, distributed across 55 contigs with a GC content of 36.59% and 5,352 protein-coding genes. Genome-genome distance and average nucleotide identity confirmed its designation as a novel strain within *P. filamentosa.* Assignment of genes/proteins in diverse functions such as drought tolerance, plant growth promotion (PGP), lantibiotics, polyketides, vitamin synthesis, siderophores production, and phosphate solubilization highlight its potential utilization in agriculture as a PGPR and industrial production of antimicrobial agent, vitamins, and biopolymers. Our research concluded that the fluorescence exhibited by JURBA-X is potentially attributed to the production of resistomycin, which might have been horizontally transferred from *Streptomyces resistomycificus* as inferred by the homology of resistomycin (rem) biosynthesis cluster genes.

## Introduction

Plants live in a cooperative environment assisted by and assisting several other organisms, including bacteria. Some of these bacteria, associated with plant roots, can ameliorate plant growth and are known as plant growth-promoting rhizobacteria (PGPR). PGPR colonizes the rhizosphere and enhances soil health, quality, and plant growth via different mechanisms. PGPR enhances plant growth promotion (PGP) by producing plant growth regulators (indole-like compounds, cytokine, and ethylene); mineral solubilization (specially for phosphate, zinc, potassium); assimilation of nitrogen and sulfur; iron sequestration (siderophores); and production of volatile compounds (2,3-butanediol and acetoin); synthesis of polyamines (spermidine, putrescine) (Grover et al., 2021). PGPR also enhances biotic and abiotic stress tolerance via induction of systemic resistance, production of antibiotics, and stress hormone ethylene by ACC-deaminase, enabling their utilization as biofertilizers and biocontrol agents (Mellidou and Karamanoli, 2022).

Among the kingdom bacteria, the PGPR belonging to the *Bacillus* genus are considered potent PGPR due to their adaptability and persistence under adverse conditions. *Bacillus* species such as *B. megaterium*, *B. thuringinesis* KR1, *B. amyloliquefaciens* p16 and *B. paramycoides* have been used as PGPR (Grover et al., 2021). In the present study, we are focusing on one strain belonging to *B. filamentosus*, recently named *Priestia filamentosa* (Gupta et al., 2020). The first report of *B. filamentosus* was a sample of marine sediments in Goa, India (Sonalkar et al., 2015). They were identified as strictly aerobic, nonmotile, Gram’s positive, and endospore-forming bacteria. *B. filamentosus, Brevibacterium casei,* and *Janibacter indicus* isolated from the rhizosphere of *Zygophyllum coccineum* showed ACC-deaminase activity, whereas the inoculation of wheat with these PGPR shows increased halotolerance (Khalifa et al., 2020).

Yahaghi et al., reported inoculation of *B. filamentosus* strain YSP110 to alfalfa plants showed 18.0 and 72.4 % of Pb accumulation in root and shoot (Yahaghi et al., 2019). Further, they also reported increased root and shoot growth of alfalfa seedlings inoculated with *B. filamentosus* strain YSP110 under a medium enriched with Zn. These data suggest *B. filamentosus* strain YSP110 is potent PGPR and microbe-assisted phytoextraction of heavy metals. Two plasmids in *B. filamentosus* HL2RP6 and *Oceanobacillus* HL2RP6 encode genes for osmoregulation. Genes such as trehalose, ectoine synthetase, alanine, proline, porins, inorganic ion transporters, peptidases and dehydrogenases play a vital role in the osmoregulation of these isolates (Mukhtar et al., 2019). Earlier, *B. subtilis* (K7), *B. filamentosus* (K1) and *Lysinibacillus cresolivorans* (K5) were isolated from desert soil samples of Cholistan, Pakistan and several thermophilic proteases isolated from *B. filamentosus* (K1) show optimum temp at 65°C, whereas these proteases can de-stain the blood from cotton cloth and de-hair the cow’s skin (Cresolivorans et al., 2021). The monoculture and consortium of *B. filamentosus, B. firmus* and *B. subterraneus* show decolorization of Novacron Red textile dye (Guembri et al., 2021). These studies show the use of *B. filamentosus*, the constitution of this consortia, and its potential application in textile water treatment.

Co-inoculation of *B. filamentosus* enhanced antioxidant status in vitamin-C fermentation (Zhang and Lyu, 2022). Magyarosy et al., first identified the Gram-positive *Bacillus* spp. under ultraviolet light. Further studies showed chlorinated resistomycin derivatives responsible for yellow fluorescence (Magyarosy et al., 2002a). In the present study, a drought-tolerant, filamentous, and fluorescent *P. filamentosa* strain was isolated from rhizosphere soil samples, for which the whole genome was assembled, followed by the identification of responsible genes for PGP traits, osmotic regulation, fluorescence activity, and antimicrobial activity.

## Materials and Methods

### Isolation and characterization of the bacteria

Rhizosphere soil samples were collected from groundnut fields in drought-hit areas of Anantapur district of Andhra Pradesh, India (location coordinates 14.1417 N 77.9831 E). Rhizosphere soil (0.5 gram) sample was dissolved in sterile saline followed by agitation for 20-30 min. The soil suspension was heated for 10 min at 70 ℃, followed by serial dilutions. 10^-3^ and 10^-4^ dilutions were inoculated on the nutrient agar (NA) medium and incubated at 28±2 °C for 24 - 48 h. Colonies with varying colors and fluorescence activity were selected for further studies. Phenotypic characters such as colony shape, edge elevation, surface motility, cell shape, Gram staining, and endospore formation were analyzed. A specific isolate with characteristics such as reddish yellow color, soft and slimy morphology, and evenly spread as a film on the agar surface (**Figure-1**) was chosen for further studies, and this isolate was named JURBA-X.

**Figure-1:**
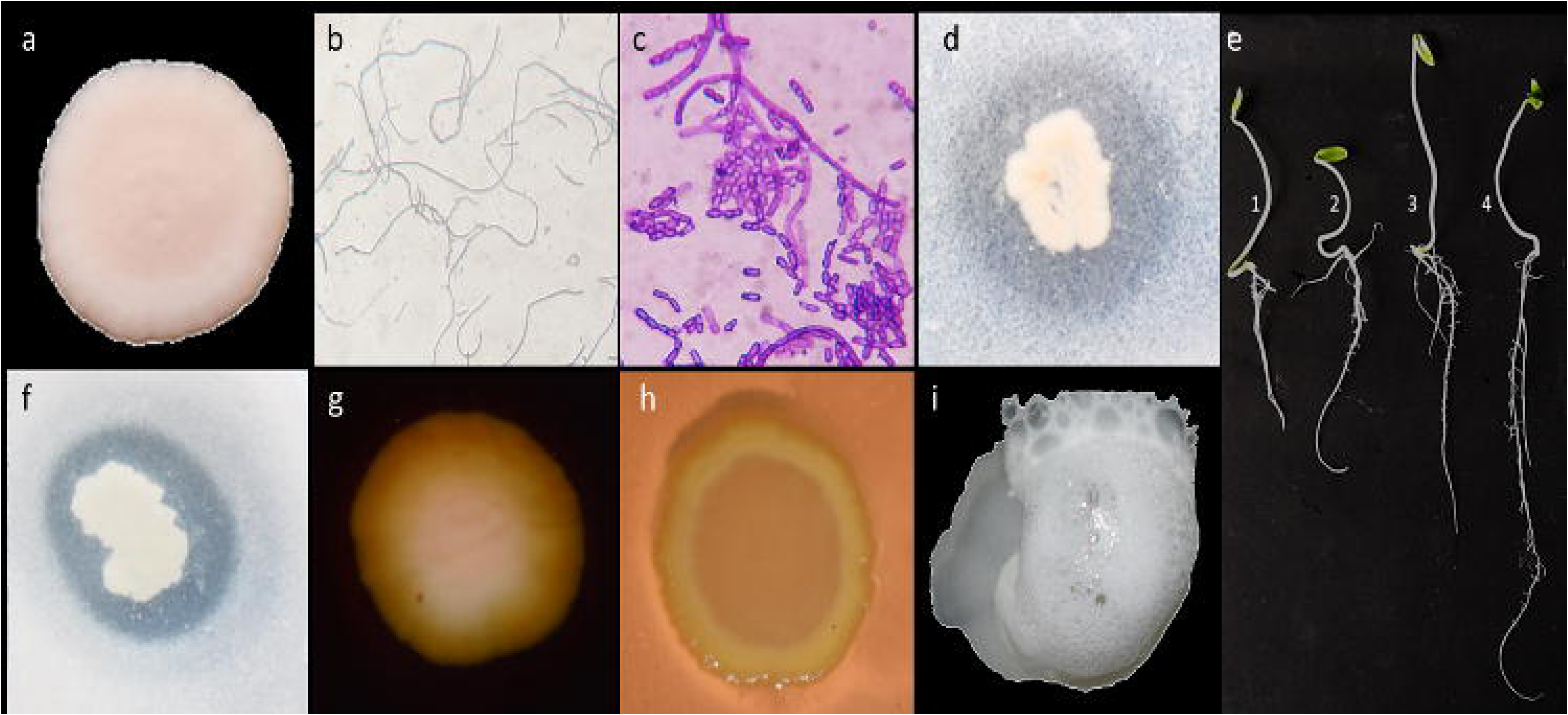
Morphology and selected enzymatic activities of *P. filamentosa* JURBA-X: (**a)** Colony morphology, (**b)** cellular morphology (under 100X scale), (**c)** Gram’s reaction (**d)** protease activity, (**e)** root development assay (cucumber seedlings), (**f)** phosphate solubilization, (**g)** starch amylase activity, (**h)** pectinase activity and (**i)** catalase activity.

### Isolation and purification of JURBA-X genomic DNA

Whole genomic DNA was isolated using the CTAB method followed by RNaseA treatment. In this process, pure culture of JURBA-X isolate was grown in 25 ml nutrient broth at 37°C overnight and 120 rpm on a shaking incubator. 15 ml of the culture was centrifuged at 12,000 rpm for 20 min at 4°C to sediment the bacterial pellet. Standard protocol was used for DNA extraction with the CTAB lysis buffer containing 2% w/v CTAB (Sigma-Aldrich, Poole, UK), 100 mM Tris–HCl (pH = 8.0; Fisher), 20 mM EDTA (pH = 8.0; Fisher) and 1.4 M NaCl (Fisher). The final DNA pellet was air-dried and re-suspended in 200 μL of 75 mM TE buffer (pH = 8.0; Sigma-Aldrich). The 260/280 ratio was identified to be >1.8, as observed by the NanoDrop ND-1000 spectrophotometer, and the integrity of DNA was confirmed to be intact using 0.8 % agarose gel electrophoresis.

### Microbial identification

16S rRNA sequencing of the isolated DNA was performed using universal bacterial primers at Eurofins Genomics India Pvt Ltd, Bangalore, India. Briefly, polymerase chain reaction (PCR) was performed using 16S rRNA gene forward and reverse primers (5’-AGAGTTTGATCCTGGCTCAG-3’ & 5’-ACGGCTACCTTGTTACGACTT-3’). The PCR amplification reaction followed a T_m_ of 52^0^C and 35 cycles of amplification of 16s rRNA gene in a 25 µl reaction mixture. PCR amplified gene product was analyzed in 1% agarose gel electrophoresis and sequenced. Homologous gene sequences of partial 16S rRNA and their respective taxonomy were identified via BLASTn against the 16S rRNA database available in NCBI. Later, after genome assembly and annotation, we also classified and confirmed its taxonomy using full-length 16S rRNA and other housekeeping gene homology and phylogeny.

### Genome sequencing

The genome of JURBA-X was sequenced with Illumina NextSeq500 platform using mate-pair (MP) and paired-end (PE) sequencing. The library profile of PE and MP were analyzed for quality and quantity using Agilent Tope station using sensitivity D1000 Screen Tope. The mean peak size and concentration of the PE library profile were 528 bp with 687 pg/µl, whereas 634 bp with 969 pg/µl for MP. PE and MP clusters were generated and sequenced using Illumina NexSeq500 2X150 bp chemistry. The MP sequencing generated 2,109,046 raw reads with ∼619 Mb data (618,597,176 bases), whereas PE generated ∼638 Mb (638,104,791 bases) data from 2,173,257 reads.

### Genome assembly, contig ordering, and scaffolding

The raw data was checked for quality confirmation using FastQC (v0.11.8). Several quality improvement strategies, such as the removal of low-quality reads (Phred Score <30) and adapter sequences, were implemented using Trimmomatic (Bolger et al., 2014) (v0.39) to ensure that only quality reads were used during further analyses. These quality reads were used to perform the *de novo* assembly using the assembly module in Tychus (Dean et al., 2018) (NEXTFLOW∼v0.23.0). This tool could construct a super (consensus) assembly of higher quality and contiguity, producing increased confidence and a comprehensive representation of the bacterial genome. Additionally, assembly improvement tools SSPACE (Boetzer et al., 2011) (v3.0) and GapFiller (Nadalin et al., 2012) (v1-10) were utilized to extend the contigs into scaffolds. Using the unmapped reads during the contig assembly step, these tools carefully detected reliable overlaps, clustered similar reads, and reconstructed the gaps to yield scaffolds. Assembled scaffolds were evaluated with QUAST (Gurevich et al., 2013) (v4.5) to assess common assembly score metrics, such as the number of contigs, largest contig, and N50.

To achieve chromosome-ordered assemblies, the JURBA-X assembly was aligned with the closest known reference genome (*P. filamentosa* Hbe603: GCA_000972245.3) to order the contigs according to the chromosomal structure as indicated in the reference. For this, we employed a reference-guided approach, using RaGOO (Alonge et al., n.d.) (v1.1), a contig ordering and orienting tool that enumerates the speed and sensitivity of Minimap2 (Li, 2018) (v2.13), to accurately achieve complete chromosome level assembly. Additionally, scaffolds mapping to the plasmids were confirmed using PLSDB (Galata et al., 2019) (v2020_03_04), a plasmid repository for bacterial plasmids, using blastn (v2.2.31). This Whole Genome Shotgun project has been deposited at DDBJ/ENA/GenBank under the accession JAGKQT000000000. The version described in this paper is version JAGKQT010000000.

### Average nucleotide identity, genome-genome distance, and phylogenetic analysis

The *Bacillus* genus comprises more than 1000 genomes. The genus *Bacillus*, harboring >293 species/subspecies, constitutes a phylogenetically incoherent group. Based on phylogenetic and molecular evidence, a recent report (Gupta et al., 2020) has robustly demarcated *Priestia* as a separate genus earlier belonging to *Bacillus*. They proposed that 17 *Bacillus* species should be recognized as a novel genus *Priestia*. To efficiently plot the phylogenetic position of our studied organism, i.e., *P. filamentosa* in the Bacillaceae family, we extracted the homologous sequences of our gene of interest using BLASTP (McGinnis and Madden, 2004) (v2.2.31) against the NR database. Based on prior literature studies (Patel and Gupta, 2020), the protein sequence of gyrase subunit B was selected as a phylogenetic marker to establish the phylogeny. The extracted sequences were aligned using Muscle (Edgar, 2004) (v3.8.1551), and the phylogenetic inference was drawn using the Maximum likelihood [ML] approach based on the JTT+GAMMA model in RAxML (Stamatakis, 2014) (v8.2.12). MEGA (Kumar et al., 2008) software was utilized to select the best phylogenetic model using the alignment under study. The phylogenetic tree was visualized using iTOL (Letunic and Bork, 2021), and information related to all neighboring clades and species were mapped to the tree.

Along with protein-marker gene-based phylogeny, we used genome-based strategies to find the close-related organisms for the organism under study. First, we calculated the genome-to-genome distances between *P. filamentosa* JURBA-X and other *Priestia* genomes (*P. filamentosa* DSM 27955, *P. filamentosa* SGD-14, *P. filamentosa* Hbe603, *P. filamentosa* PK5_39, *P. filamentosa* 2102, *P. filamentosa*, *P. filamentosa* HL2HP6, *P. endophyticus*, *P. megaterium*, and *P. koreensis* using GGDC tool (Auch et al., 2010a) (version 2.0 and Formula 2). We also calculated the average nucleotide identity (ANI) between the abovementioned organisms using the pyANI algorithm (Pritchard et al., 2016).

### Comparative genomics

For comparative analysis, five closely related *Priestia* genomes viz., *P. filamentosa Hbe603*, *P. filamentosa 2102*, *P. endophytica*, *P. koreensis*, and *P. megaterium* were selected, and their information was downloaded from NCBI. CGView (Grant and Stothard, 2008) server was utilized to generate comparative maps against the reference to *P. filamentosa* JURBA-X, using BLAST homology comparisons and was represented as a circular genome plot in **Figure-2**.

**Figure-2:**
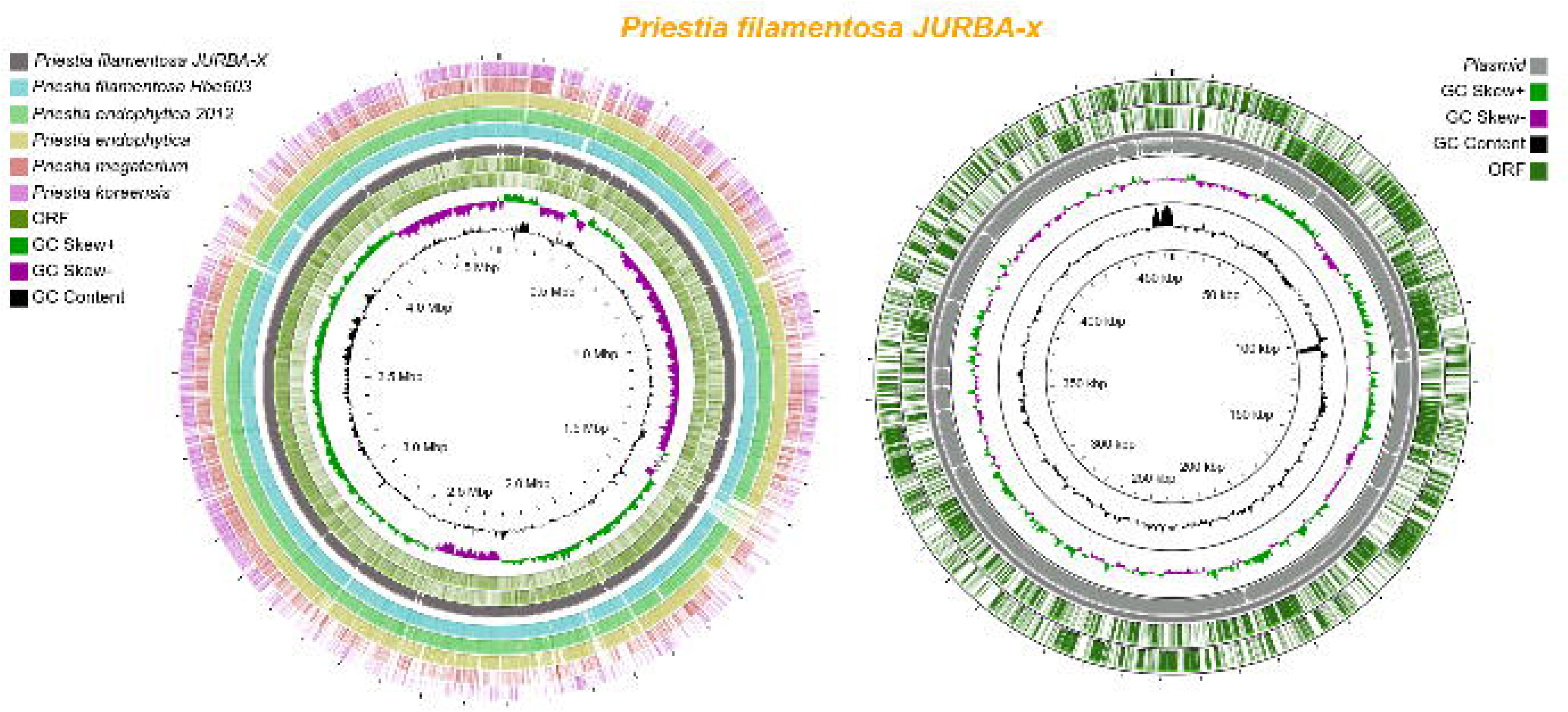
Circular representation of *P. filamentosa* JURBA-X chromosomal and plasmid genome assembly. Circles (from inside to outside) 1 and 2 (GC content: black line and GC skew: magenta and green lines), circle 3 (*P. filamentosa* JURBA-X: red circle); circle 4 (mapped *P. filamentosa* Hbe603 genome with *P. filamentosa* JURBA-X genome: green circle); circle 5 (mapped *B. endophyticus* 2102 genome with *P. filamentosa* JURBA-X genome: purple circle); circle 6 (mapped *P.* endophytica genome with *P.* filamentosa JURBA-X genome: Orange circle); circle 7 (mapped *P.* megaterium genome with *P.* filamentosa JURBA-X genome: blue circle); circle 8 (mapped *P.* koreensis genome with *P.* filamentosa JURBA-X genome: yellow circle). Mapping studies were performed using BLASTn and CGView server was used to build the circular representation.

### Genome annotation and functional characterization

Gene prediction and functional annotation were performed using Rapid Annotation using Subsystem Technology (RAST) (Aziz et al., 2008) (v2.0). Further, annotated protein sequences were subjected to different functional characterization databases to identify the key functions, pathways, ontology, and more. KEGG Automatic Annotation Server (KAAS) (Moriya et al., 2007) was used to carry out the functional pathway analysis where the total proteins were subjected to the bi-directional best hit (BBH) method assigning orthologs. For GO analysis, HMMER2GO (Finn et al., 2011) (v0.17.9) was used against a customized HMM Pfam (Finn et al., 2014) database to obtain information on the molecular functions, cellular components, and biological processes associated with the proteins and the results were visualized in WEGO (Ye et al., 2018) (v2.0). COG categories were identified within the *P. filamentosa* JURBA-X genome by subjecting the proteome to rpsBLAST (v2.2.31) against NCBI’s Conserved Domain Database (CDD) (Marchler-Bauer et al., 2011) at E-value 1e-2. Subsequently, COG categories were assigned to query protein sequences using COG phylogenetic classification. To identify functional domains present in each annotated protein, all protein sequences were subjected to hmmscan (Finn et al., 2011) (v3.1b2) against the PFAM-A database (Finn et al., 2014), followed by parsing of the data and protein architecture generation. The characterized GO and COG categories, PFAM domains, and KEGG pathways related to diverse biological processes were visualized via Bar Plots and Pie charts (R-4.0.2).

### Secondary metabolites identification

BAGEL4 (Van Heel et al., 2018) was used for identifying the presence of bacteriocins. Using genome fasta file as an input, it identifies lantibiotics and peptides. Metabolites were characterized for potential solutions in the JURBA-X genome. With the aid of literature (Magyarosy et al., 2002b) we also utilized the antibiotics and secondary metabolites analysis shell (antiSMASH) (Blin et al., 2019) tool for the identification of all secondary metabolites. The presence of fluorescent compounds in the isolate JURBA-X was also validated using mass spectroscopy analysis, where the extracted compound’s spectroscopic values were similar to chloroxanthomycin / resistomycin (REM). We also identified type II polyketide synthase metabolite and its complete REM gene cluster in the JURBA-X genome using blastp. For this analysis, we created a custom database of 18 REM cluster proteins as identified in the previous study (Jakobi and Hertweck, 2004a) and further used BLASTP (McGinnis and Madden, 2004) (v2.2.31) to substantiate the presence of REM proteins in the JURBA-X genome. REM proteins that were identified using NR Diamond Blast and ANTISMASH (Blin et al., 2019) prediction were mapped against *Streptomyces resistomycificus* REM gene cluster using GGgenes (R-4.0.2, v0.4.1) to understand the organization of the resistomycin (rem) biosynthesis gene cluster within JURBA-X. Later, each of the core REM proteins, i.e., RemA, RemB, RemC, RemF, RemI, and RemL were used as phylogenetic markers to recognize closely related homologous genomes in the Bacillaceae family with the presence of REM cluster indicating fluorescence. MEGA (Kumar et al., 2008) (v2.2.4) software was used to carry out phylogenetic tree reconstruction, and ITOL (Letunic and Bork, 2021) was used for visualization of the tree.

## Results and discussion

### Morphological characteristics of JURBA-X

Strain JURBA-X can be morphologically classified and identified (**Figure 1**) as a filamentous, Gram-positive, and endospore-forming bacterial isolate (**Figure 1a, b, c**). The colonies are mild pink/peach pigmented, circular, granular dry colonies with 1.3 ± 7 mm diameter. Colonies grown on NA or TSA show strong yellow-orange fluorescence under UV illumination. Cells are nonmotile and tolerant to osmotic stress (-0.73 MPa). JURBA-X shows positive outcomes for synthesizing siderophore, protease (**Figure 1d**), exopolysaccharides, phosphate solubilization (**Figure 1f**), and incompatibility between the Gram-positive and negative bacterial isolates. Our results further identified the ability of this strain to produce indole-like compounds, chitinase, amylase (**Figure 1g**), cellulase, and pectinase (**Figure 1h**) and show catalase activity (**Figure 1i**). JURBA-X strain is sensitive to antibiotics chloramphenicol (30 µg), Ciproflaxin (5 µg), tetracycline (30 µg), and vancomycin (30 µg) and shows an average of 2.8 cm, 4.2 cm, 4.2 cm, and 2.6 cm zone of inhibition around the colony, respectively. Cucumber seeds primed with JURBA-X showed no significant seed germination efficiency and showed increased root growth development (**Figure 1e**).

Pigmentation and fluorescence activity are restricted to growth on solid media, whereas no detectable fluorescence activity was observed in the liquid media. The culture medium was supplemented with essential amino acids, trace elements (Cu, Zn, Mn), and sugars (dextrose, fructose, arabinose, and lactose). Changes in culture conditions, such as the presence and absence of light, pH (5-9), and temperature (0-50°C), caused no effect on the pigmentation and fluorescence activities. Solvents, i.e., methanol, ethyl acetate, isopropanol, diethyl ether, etc., were ineffective for extracting the fluorescent compound without cell wall lysis. Tween-20 (0.05%) in methanol followed by cell-wall lysis proved an effective method for extracting the yellow fluorescent compound.

### Genome assembly and annotation of JURBA-X

Illumina NextSeq500 paired-end (PE) technology generated 2,173,257 reads of 100-151 bp each, accumulating 638,104,791 bp. *De novo* assembly of 2,106,788 high-quality reads resulted in a draft genome of 5,113,908 bp length distributed in 55 contigs, with the largest contig length of 1,246,062 bp and average GC content of 36.42%. With the help of reference-based contig ordering against available genomes having chromosomes and plasmids, 24 contigs were assigned to the chromosome and 31 to the plasmid. While the largest contig of 1,246,062 bp belonged to the chromosome, a contig of 112,570 bp with an average GC content of 34.81% was the longest contig belonging to the plasmid. The N50 value of chromosome and plasmid was found to be 499,773 bp and 50,978 bp, respectively.

RAST-based annotation identified 5,004 putative coding regions within the chromosome assembly, of which 3,653 proteins (73.00%) were functionally annotated, while the remaining (23.00%) were hypothetical proteins. The remaining coding regions were characterized as RNAs [0.65%] or tRNAs [3.71%]. Similarly, RAST identified 600 putative coding regions within the plasmid, of which 263 proteins (43.83%) were functionally annotated, while the remaining (51%) were hypothetical proteins. The rest of the coding regions were characterized either as RNAs [1.00%] or as tRNAs [4.5%]. Genome assembly and annotations statistics have been tabulated in **Table-1**, and the annotated genes, GC skew, and GC content across the genome can be visualized in **Figure-2**.

**Table 1.**
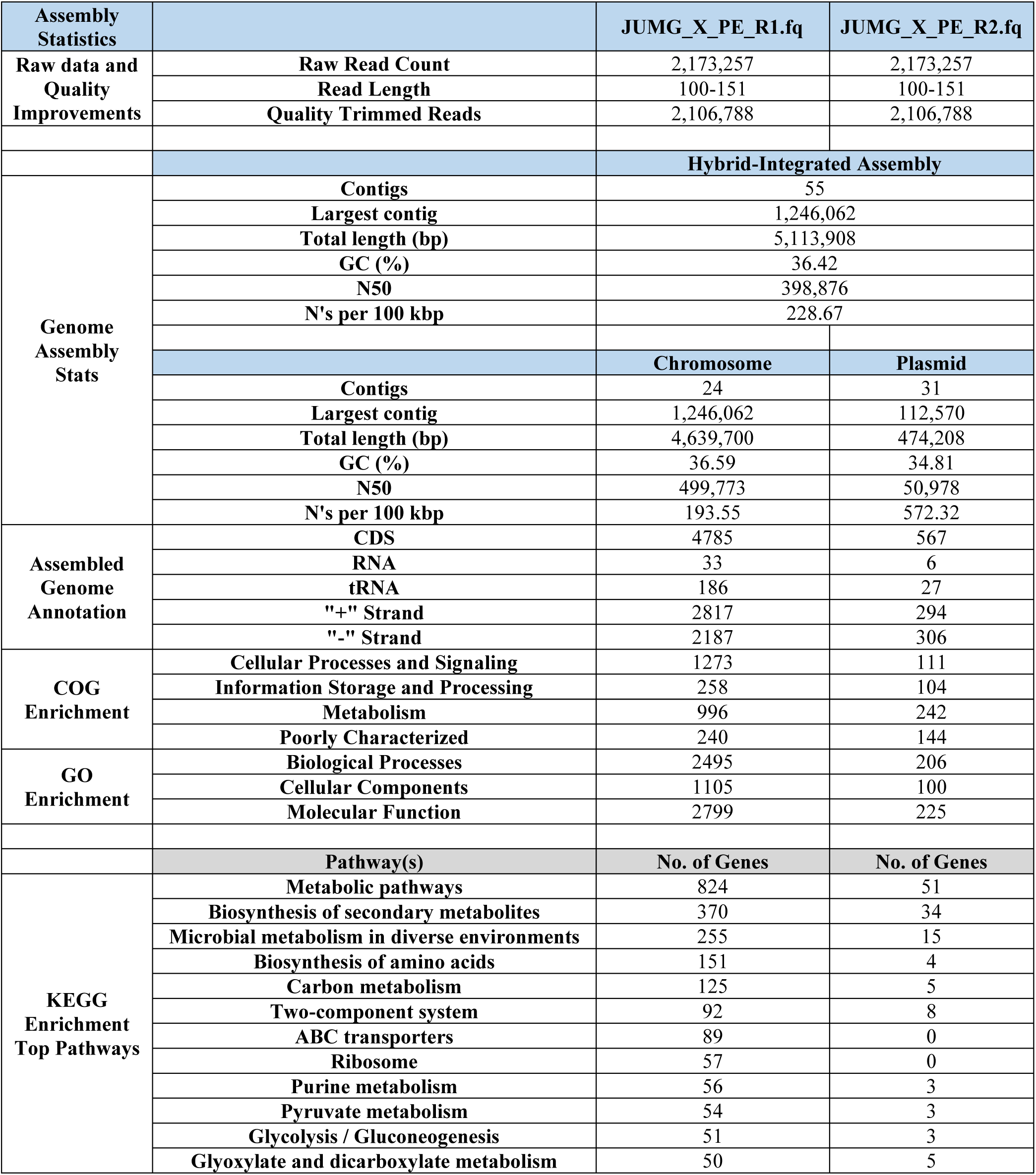
Assembly statistics for *P. filamentosa* JURBA-X chromosome and plasmid sequence.

### Phylogenetic analysis of the *Priestia filamentosa* clade

Based on 16s rRNA sequencing and molecular phylogeny, JURBA-X has been identified as a member of *P. filamentosa.* The members of genus *Priestia* were previously associated with *Bacillus*, known as Gram positive bacteria with rod shape. The members of *Priestia* are predominantly present in diverse locations such as cotton inner tissue, sea sediments and willow plant rhizosphere. The genus *Bacillus* comprises more than 1000 genomes, harboring 293 species/subspecies, constituting a phylogenetically incoherent group. To be efficiently able to plot the phylogenetic position of *P. filamentosa* within the Bacillaceae family is a challenging task and does not have an easy resolution. The taxonomic position of *P. filamentosa* JURBA-X was tried to resolve based on phylogeny of 16S rRNA and one of the known phylogenetic marker DNA gyrase B (GyrB), average nucleotide identity, and genome-genome distance.

The preliminary phylogenetic tree using 16S rRNA marker sequences was not able to resolve all species of the *Priestia* genus (**Figure-S1**). *P. filamentosa* strain SGD-14, *P. endophyticus* strain 2DT, *B. kochii* strain WCC 4582, *B. purgationiresistens* strain DS22 and *B. ciccensis* strain 5L6 were found to form a group together. Despite its popularity, 16S rRNA is not a credible marker for taxonomic placement below the genus level, specifically for the organism of interest *P. filamentosa.* Therefore, housekeeping gene gyrase-B analysis was performed to validate the taxonomic relationship among *Priestia* and *Bacillus* genus.

The ML phylogenetic tree generated for gyrase-B (**Figure-3**) showed the summary of all JURBA-X neighboring clades and species in an efficient manner. All trees exhibited nearly clear branching of *Priestia*, and *Bacillus* species and consistently displayed clades encompassing *Priestia* species such as *P. endophytica, P. koreensis, P. megaterium, P. aryabhattai, P. flexa, P. abyssalis* and *B. aquamarine, B. vietnamensis* interspersed with other Bacillaceae species, adjacent to *Cytobacillus* species. *Listeria monocytogenes* was chosen as an outgroup for this study.

**Figure-3:**
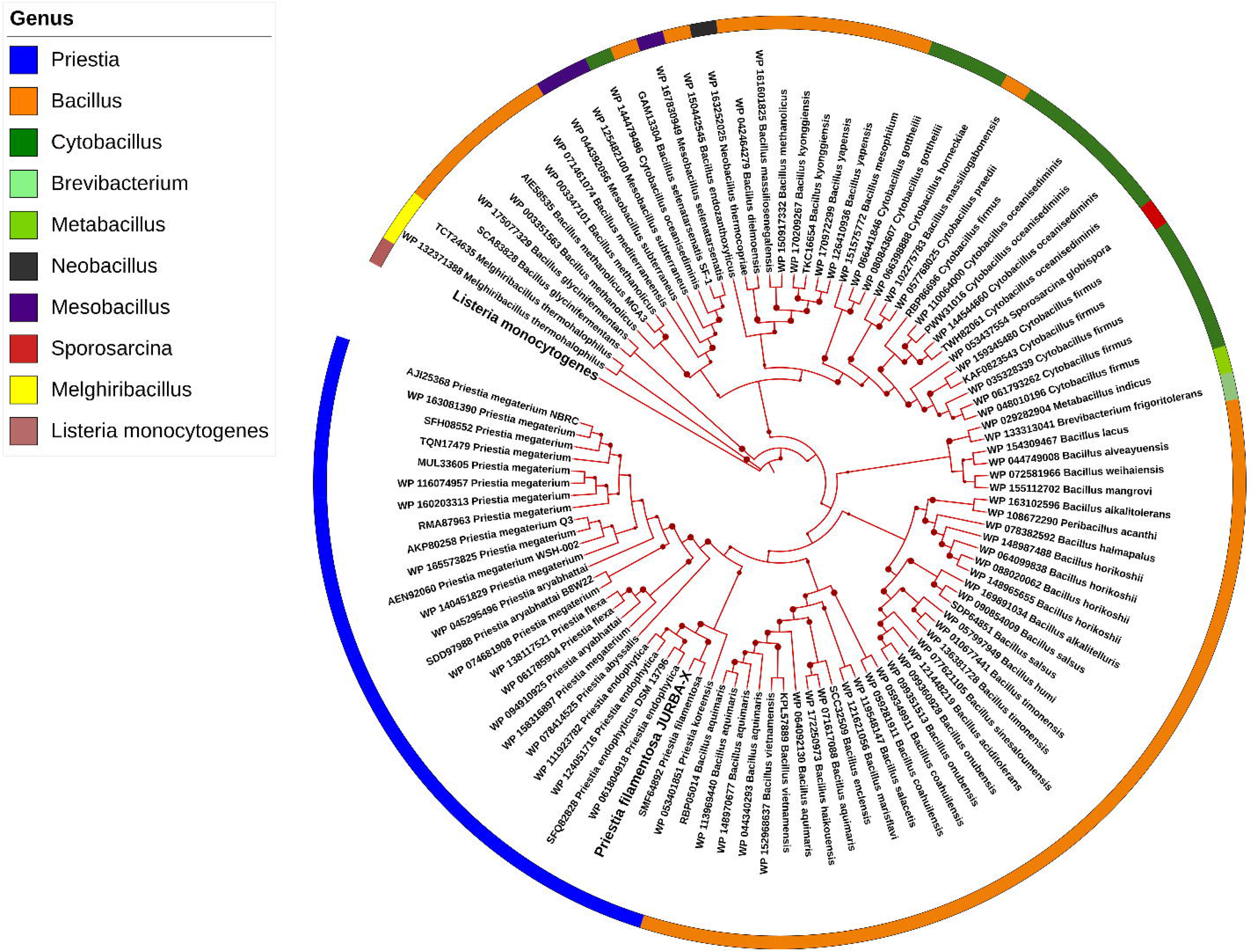
Phylogenetic characterization of *P. filamentosa* JURBA-X using a single-copy housekeeping protein i.e., DNA gyrase subunit B reveal the correct taxonomic placement of all closely related species.

Taxonomic classification using ANI resulted in inference of species level identification to *P. filamentosa* species with a perfect confidence score of 100% identity. However, ANI identity score against *P. filamentosa* Hbe603 was found to be 99.05%. This implies the Hbe603 strain of *P. filamentosa* has some divergence to JURBA-X, but they display high nucleotide identity because they both belong to the same species i.e., *P. filamentosa*. ANI identity scores against species *P. endophytica, P. megaterium,* and *P. koreensis* and some other strains have been tabulated in **Table-S1**. This analysis clearly infers that these are all closely related species but have definite demarcation to be assigned as a separate taxonomic classification.

We also estimated DNA-DNA hybridization (DDH) values between genus *Priestia* and *Bacillus* members using GGDC (Auch et al., 2010b) (Genome-To-Genome Distance Calculator) server which uses GBDP strategy (Genome Blast Distance Phylogeny). The GGDC values also jointly concurred that JURBA-X has a close genomic distance to all species and strains of *Priestia* but still has some differentially characterized genomic features making it a novel strain [**Table-S2**].

### Functional characterization of JURBA-X genome

The annotated proteomes were subjected to functional enrichment analysis to identify proteins of interest and establish association to various pathways. The ability of biologists to analyze and interpret such data relies on functional annotation of the previously known proteins. These analyses help characterize organisms thereby establishing functional evidence necessary to infer their biological relevance.

Functional characterization using the KEGG database provided insights into pathways representing the molecular interactions of JURBA-X. Other than the housekeeping metabolism pathways, a major portion of the JURBA-X genome [both chromosome and plasmids] was characterized to possess genes of “Biosynthesis of secondary metabolites” and “Microbial metabolism in diverse environments” (**Figure-4A**). This was of specific importance, since early culture-based characterization of JURBA-X showed fluorescent and antibacterial activities, which largely belonged to these two pathways. On one hand, there have been only few reports of presence of fluorescent properties within the Bacillaceae family and on the other, antibiotic properties are commonly identified across various strains of *Priestia* and *Bacillus*. For both these reasons, we proceeded with further characterization of fluorescence and antibacterial properties of JURBA-X

**Figure-4:**
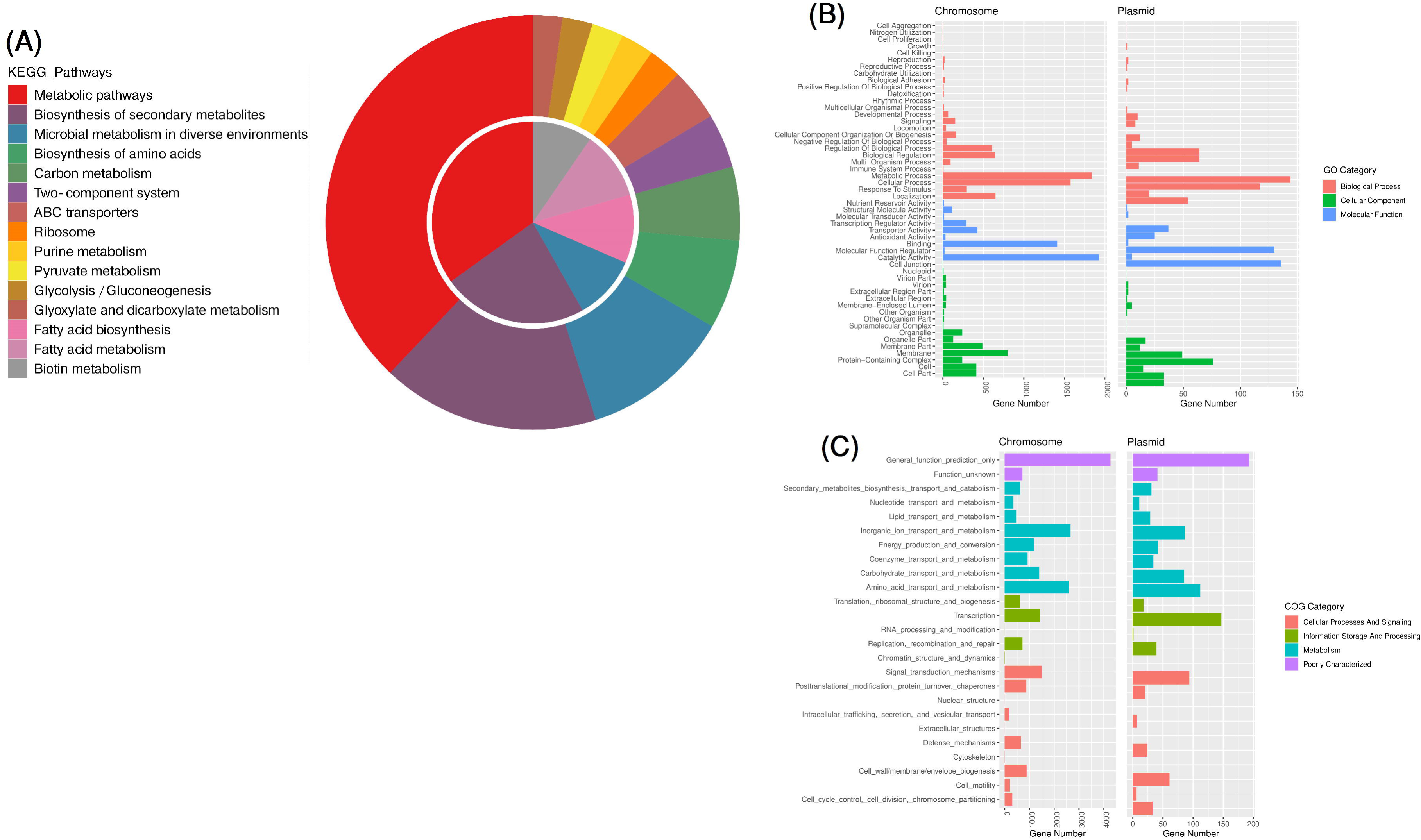
Illustrations representing the distribution of (**A)** KEGG pathways, (**B)** Gene Ontology (GO) and (**C)** Cluster of Orthologous groups (COG) categories associated with the genome of *P. filamentosa* JURBA-X as identified using KAAS - KEGG Automatic Annotation Server, GO, and COG databases.

The Gene Ontology (GO) Analysis of JURBA-X Chromosome identified a total of 2,495 of coding genes belonging to “Biological Processes”, 1,105 of them belonging to “Cellular Components” and 2,799 coding genes belonging to “Molecular Functions” of the GO database. Whereas, in the plasmid region of JURBA-X genome, 206, 100, and 225 coding genes were classified to “Biological Processes”, “Cellular Components” and “Molecular Functions” respectively (**Figure-4B**).

Amongst the total proteome of the chromosome region, ∼ 57% (2,767) proteins were categorized into Clusters of Orthologous Groups (COGs). Among these mapped proteins, ∼21% belonged to metabolism functions, ∼26% to cellular processes & signaling, and ∼ 5% belonged to information storage & processing. According to COGs mapping, 144 proteins were involved in inorganic ion transport & metabolism (COG: P), 104 proteins were involved in signal transduction mechanisms and 54 proteins were reported to function in secondary metabolites biosynthesis, transport, and catabolism (COG: Q) (**Figure-4C**). Amongst the total proteome of the plasmid region, ∼ 96% (601) proteins were categorized into Clusters of Orthologous Groups (COGs). Among these mapped proteins, ∼39% belonged to metabolism, ∼18% to cellular processes & signaling and ∼17% proteins to information storage & processing. According to COG mapping, 47 proteins were involved in inorganic ion transport and metabolism (COG: P), 40 proteins were involved in signal transduction mechanisms and 19 proteins were reported to function in secondary metabolites biosynthesis, transport, and catabolism (COG: Q) (**Figure-4C**).

### Genetic basis for osmotic regulation

#### Osmotic regulation

In plants, the adaptation to drought stress is associated with metabolic adjustments leading to the accumulation of several compatible solute/osmolytes like proline, sugars, polyamines, betaines, quaternary ammonium compounds, polyhydric alcohols, amino acids and water stress proteins like dehydrins (Yancey et al., 1982). Glycine betaine is the most potent osmolyte, distributed in a wide range of bacteria and synthesized via a two-step pathway from choline. The genome of JURBA-X contains *gbsB* gene product that can convert choline to betaine aldehyde. Along with this, *gbsA* gene product can help in synthesizing the glycine betaine from the reaction intermediate. Genome also contains gdhA, gltB, gltD, gln genes for glutamate biosynthesis, clsA/B and clsC for cardiolipin biosynthesis and specific PTS systems for trehalose, maltose, cellobiose and sucrose.

#### Antioxidant system

Stress tolerance is also associated with the expression of stress response proteins and activation of the ROS system. Genes such as *groES, groEL, dnaJ, dnaK, hslO, csaA, cop2 clpB,* and cold shock proteins (*csp*) were involved in stress tolerance. The oxidative stress response genes identified in the annotated JURBA-X genome included peroxidases - vanadium chloroperoxidase, thiol peroxidase (Txp and Bcp-type), non-heme chloroperoxidase and glutathione peroxidases (*gpx*); superoxide dismutase - Mn superoxide dismutase (Mn*sod*), Cu-Zn superoxide dismutase (Cu-Zn*sod*), Fe superoxide dismutase (Fe*sod*) and catalase - Mn catalase (Mn-*cat*) and multifunctional catalase (*cat*) as shown in **Table-S3**.

#### Spermidine

Polyamines are polycations that regulate the abscisic acid biosynthesis, scavenging of ROS, increased rate of photosynthesis, root architecture, and accumulation of osmolytes. *Arabidopsis* roots associated with the *B. megaterium* strain BOFC15 under drought stress produce extracellular polyamine spermidine enhancing plant growth and drought resistance (Zhou et al., 2016). Polyamine putrescine has also been reported to trigger disease resistance in *Arabidopsis* against bacterial pathogens (Liu et al., 2019). We identified the presence of speD (S-adenosylmethionine decarboxylase), speE, (spermidine synthase) sepA, (arginine decarboxylase) speB, (agmatinase) aguA (agmatine deiminase) and aguB (N-carbamoylputrescine amidase) genes for biosynthesis and potA (spermidine/putrescine import ATP-binding protein) for spermidine or putrescine transport system in the JURBA-X genome (**Table-S3)**.

### Genetic Basis for plant growth promotion traits

#### Siderophore biogenesis

Siderophores are small organic compounds produced by PGPR which can act as bio-control agents, and therefore allow the plant growth promotion and disease suppression of soil born phytopathogens. Genus *Bacillus* organisms produce catechol-based siderophore bacillobactin, which involves the chelation of ferric ion (Fe^3+^) from the environment and transported back into the bacterial cytoplasm via ABC transporters (Nalli et al., 2023). JURBA-X genome contains dhbA (2,3-dihydro-2,3-dihydroxybenzoate dehydrogenase), dhbB (isochorismate lyase), dhbC (isochorismate synthase), dhbE (2,3-dihydroxybenzoate-AMP ligase) and dhbF (nonribosomal peptide synthetase), necessary for bacillibactin biosynthesis. It also encodes genes for transport (siderophore ABC transporters) and import of Fe-bacillibactin (feuA, feuB, and feuC). This study revealed the presence of a few other genes such as fhuC, fhuB and fhuG, important for ferrichrome transport (**Table-S3**).

#### Phosphate metabolism

Phosphonate is an organophosphate and rich source of soil phosphate with direct carbon-phosphorus (C-P) linkage (Lo et al., 2018). Bacteria such as *P. psychrotolerans* and *B. subtilis* utilizes the phosphate-specific transport (pst) system for the free inorganic phosphate transport. Recent studies of (Kang et al., 2020) showed the pst operon structure in *P. psychrotolerans*, consisting of pstS, pstC, pstA, pstB, and two-component signal transduction system phoP/phoR genes for the uptake of inorganic phosphate. The whole-genome analysis of the JURBA-X showed pst operon (pstA, pstB, pstC, and pstS genes) and phoB, phoH, phoP, phoR and phoU genes (**Table-S3**). This suggests that JURBA-X could possibly regulate the phosphate and polyphosphate degradation via polyphosphate kinases (ppk), exopolyphosphatase (pph), and alkaline phosphatase. The bioavailability of phosphate from phosphonate is mediated by bacterial phn gene action. The genome of JURBA-X also contains genes for organophosphonate (phnC, phnD, and phnE) metabolism. Our experimental results also revealed the ability of this strain to degrade phosphate on petri plate (**Figure 1f**).

#### Volatile organic compounds

The volatile organic compound acetoin and 2,3-butanediol are released by rhizobacteria and enhance the growth of plant under drought stress and disease resistance (Russo et al., 2022). It has been reported earlier that *Arabidopsis thaliana* and *Nicotiana benthamiana* treated with acetoin and 2,3-butanediol from the *B. amyloliquefaciens* BZB42 shows the stomata closer via salicylic acid and ABA signaling pathway (Wu et al., 2018). The synthesis of acetoin is catalyzed by the ilvBGI and ilvHN (acetolactate synthase) and budA (acetolactate decarboxylase) from pyruvate to acetoin. The acetoin further converts to 2,3-butanediol either by host plant or bacteria. The JURBA-X genome contains ilvBGI and ilVHN which catalyze the conversion of pyruvate to acetoin. Therefore, the presence of these genes could be suggested to help in the plant growth promotion and systemic disease resistance.

#### Genetic Basis for antimicrobial and fluorescence activity

JURBA-X was identified as showing antagonistic activity against both Gram positive and negative bacterial isolates. The fluorescence activity was able to identify a metabolite and, TLC with methanol extract shows a single yellow fluorescence band under UV. The extract also shows antibacterial activity against *Bacillus* spp. Magyarosy et al., (2002) first reported a fluorescent *Bacillus* spp. which can produce a yellow-orange, fluorescent pigment, where the compound was identified as a halogenated derivative of resistomycin (chloroxanthomycin) (Magyarosy et al., 2002c). Resistomycin, which are type II polyketides, shows antibacterial activity against Gram positive bacteria which works as an RNA polymerase inhibitor, HIV protease inhibitor, and can induce apoptosis (Riaz et al., 2020).

On a similar note, bacteriocins are peptide antibiotics classified into two classes; class I bacteriocins are lantibiotics that undergo post-translational modifications. LanM is lantipeptide synthetase perform. The precursor propeptide is called LanA synthesized by ribosomally, contain N-terminal ladder sequence and C-terminal propeptide. Serine and threonine residues in propeptide undergo dehydration reaction and form dehydroalanine (Dha), dehydrobutyrine (Dhb) and formation of intramolecular thioether bond (lanthionine) with a cysteine residue. Single lantipeptide synthetase (LanM) catalyzes both reactions in type II lantibiotics, the mature peptide exported by ABC-type exporters (SunT). The genome of JURBA-X contains a cluster of genes for precursor propeptide LanA, LanM and SunT. The presence of gene components for resistomycin biosynthesis and lantibiotics could be a possible reason for fluorescence activity and antagonistic activity against bacteria.

#### Vitamin K

Menaquinone (MK) or fat-soluble vitamin K_2_ play an important role blood coagulation process also a key component in many bacterial electron transport system (Mladěnka et al., 2022). Here we identified *menCEBHDFA* genes cluster of MK biosynthesis and Hotdog fold thioesterase (catalyze the conversion of 1,4-dihydroxy-2-naphthoyl-coenzyme A (DHNA-CoA) to 1,4-dihydroxy-2-naphthoate) and menH which catalyze the final reaction in MK biosynthesis present in other loci of the genome.

### Antibacterial Activity of JURBA-X

Bacteriocins (Simons et al., 2020), which are largely produced by Gram-Positive bacteria, have been classified into 4 classes. Class I or lantibiotics includes small sized (<5 kDa) and post-transcriptionally modified bacteriocins. Class II or non-lantibiotics, do not contain unusual amino acids in their structure and post-translational modification is limited to bisulfide bridge formation in only a few members. Class III bacteriocins are large heat-labile lytic or non-lytic peptides (>30 kDa). Class IV bacteriocins are specified by their structure, containing lipid or carbohydrate parts (Simons et al., 2020). BAGEL4 (Van Heel et al., 2018) analysis [**Table-S4**] showed the array of bacteriocins identified across *Priestia* and *Bacillus* strains. Of these, JURBA-X was identified with class II bacteriocins only. Antimicrobial lasso peptides and LAPs have been identified in JURBA-X along with some other strains of *Priestia* like Hbe603. BAGEL4 also characterized the presence of a putative lantibiotic biosynthesis protein within the JURBA-X genome. Sonorensin, an antimicrobial peptide of class II belonging to the heterocyclic anthracin subfamily of bacteriocins was also identified to be present in JURBA-X. Hence, we concur with the presence of antibacterial compounds contributing to antimicrobial antagonistic nature of JURBA-X.

### Secondary Metabolites identification followed by the characterization and evolutionary exploration of fluorescent compound

Many microbial genomes contain several (up to 30-40) gene clusters encoding for biosynthesis of secondary metabolites. Thus, mining genetic data has become a particularly important method in modern genome screening approaches for bioactive compounds, like antibiotics or chemotherapeutics.

In the history of the Bacillaceae family there have been few reports of the presence of fluorescent compounds, which contributes to many novel properties of the organisms. Bright fluorescent orange pigments of *B. endophyticus* AVP-9 (Kf527823), with absorption maxima at 493 nm, have indicated that the pigment shows characteristics of carotenoids (Ram et al., 2017). A Gram-positive *Bacillus sp.* that fluoresces yellow, under long-wavelength UV light on several common culture media, was isolated from soil samples and reported to be chlorxanthomycin in December 2020 (Magyarosy et al., 2002b). Blue fluorescence of endolithic *Bacillus* isolates, from granite, was also reported to have fluorescent properties (Fajardo-Cavazos and Nicholson, 2006). Taking a cue from here, we investigated for the presence of similar fluorescence compounds in JURBA-X.

The light pink fluorescence of JURBA-X colonies (**Figure-5**) and the mass spectroscopy values of the extracted components strongly indicated presence of chloroxanthomycin / resistomycin. This was first validated with the identification and characterization of rem biosynthetic gene cluster within the JURBA-X genome. The enrichment of pathway belonging to biosynthesis of secondary metabolites via KEGG pathway analysis (**Figure-4A**), and the confirmation of the presence of type II polyketide synthase (T2PKS) via AntiSMASH (Blin et al., 2019) (v5.0), further validated the presence of fluorescent function within JURBA-X. Furthermore, a comparative study across several strains of *P. filamentosa* [**Table-S5**], predicted that T2PKS uniquely belonged to JURBA-X chromosome and plasmid regions.

**Figure-5:**
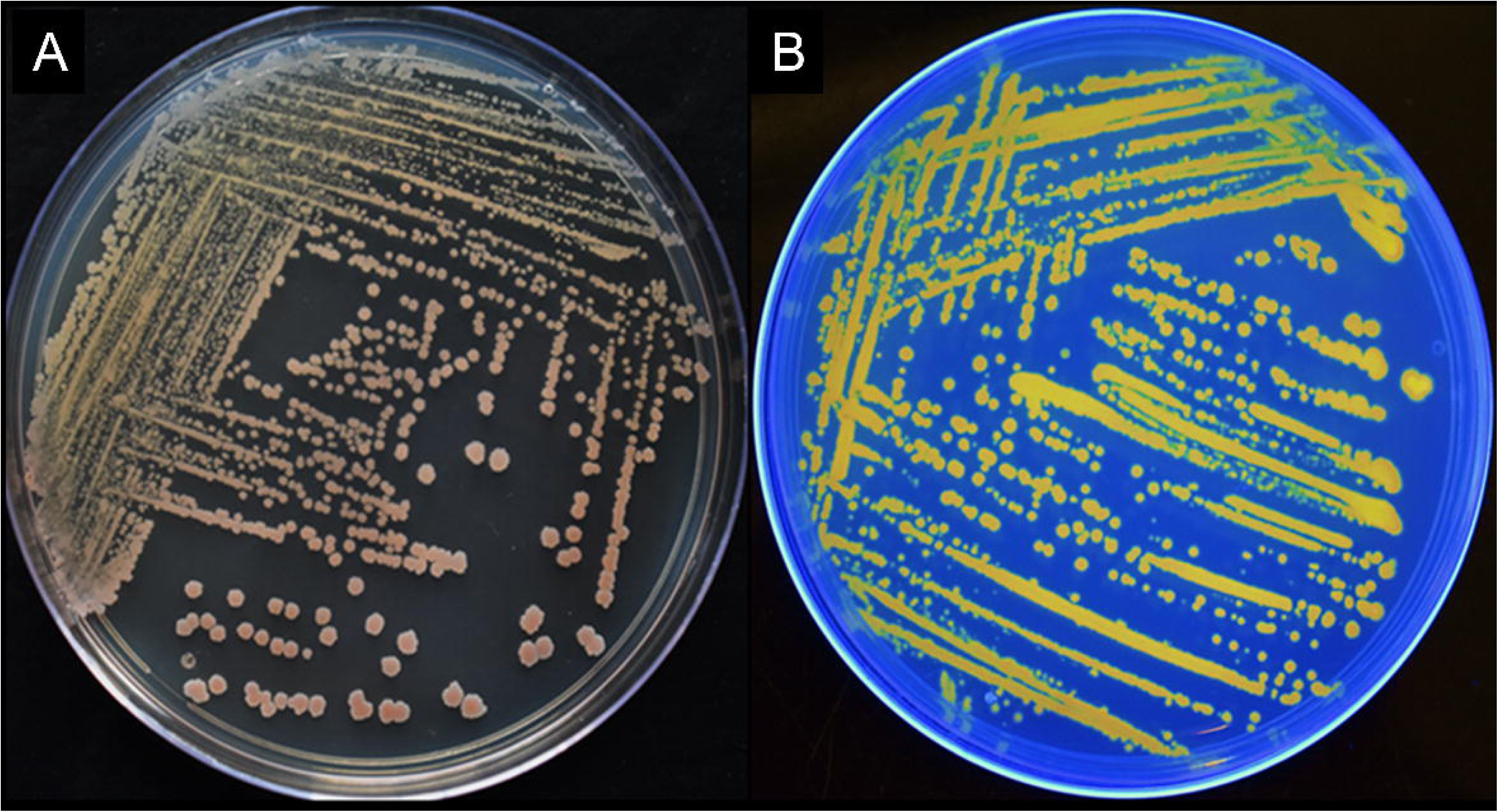
(A) Pure culture of JURBA-X with mild pigmentation and (**B)** strong florescence activity of JURBA-X pure culture grown on NA medium.

*Streptomycetes* synthesize aromatic polyketides by type II polyketide synthases (PKS), a complex of iteratively used individual proteins. Resistomycin (rem) is an unusual aromatic polyketide metabolite of *Streptomyces resistomycificus* that exhibits a variety of pharmacologically relevant properties, such as, inhibitor of HIV-1 protease, RNA polymerase and DNA polymerase, and activity against Gram-positive bacteria and mycobacteria (Jakobi and Hertweck, 2004b). Frame analysis of the 15 kb rem biosynthesis gene cluster (GenBank accession nr. AJ585192), which is flanked by genes involved in primary metabolism and housekeeping, revealed 18 open reading frames (ORFs) that consist of 11 structural genes and 7 genes for regulation and resistance (Fajardo-Cavazos and Nicholson, 2006).

Upon close examination of the T2PKS region, it was understood that the JURBA-X plasmids might have the rem (Resistomycin) cluster. Using BLASTP (McGinnis and Madden, 2004), it was determined and verified that the JURBA-X plasmids contain 11 of the 18 REM proteins known to form the rem cluster. This rem cluster organization and synteny was compared across *Streptomyces resistomycificus* (Jakobi and Hertweck, 2004a) to clearly characterize the presence of this aromatic polyketide in the plasmid as depicted in **Figure-6**. The core REM proteins, i.e., RemA, RemB, RemC, RemF, RemI, and RemL (Silvennoinen et al., 2009) were further used as phylogenetic markers to recognize other closely related homologous genomes in the Bacillaceae family, and to also explore the presence of REM cluster amongst them indicating the presence of fluorescence activity. It was observed that no other organism from the Bacillaceae family shows the presence of the REM proteins except for *B. cereus* and *P. endophytica*. It was also inferred from the phylogenetic analysis that the REM proteins are present most commonly across the Terrabacteria clade which are the common ancestors for phylum Firmicutes, Chloroflexi, and Actinobacteria as depicted in **Figure-7** and **Figure S2**. The rest of the tree was populated with *Streptomyces* spp., which is the original source of rem cluster.

**Figure-6:**
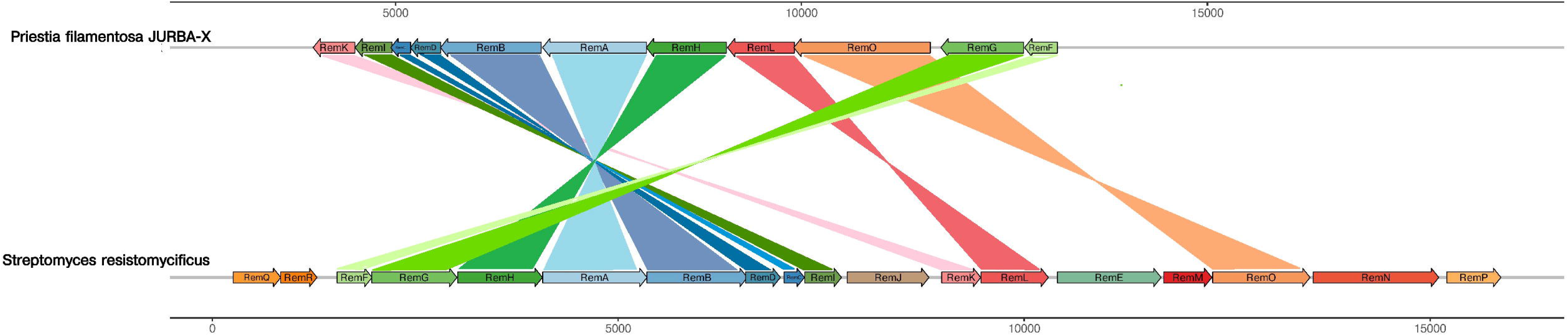
Comparative genomic organization of the resistomycin (rem) biosynthesis gene clusters between *P. filamentosa* JURBA-X and *Streptomyces resistomycificus* revealing their close synteny and distribution.

**Figure-7:**
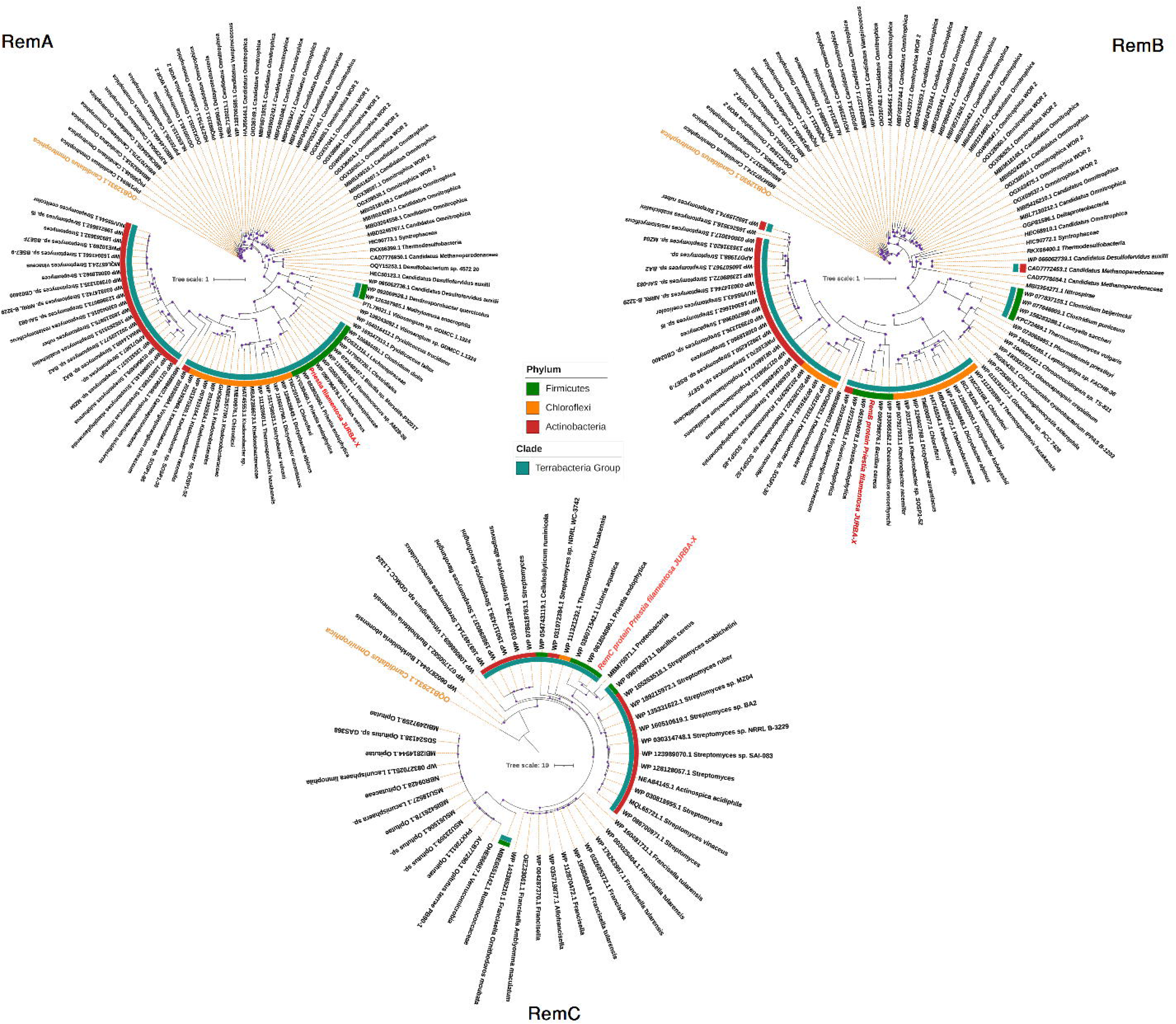
Phylogenetic studies of (a) RemA, (b) RemB and (c) RemC proteins revealing their close homology with Actinobacteria spp. and confirming the evolution of rem cluster from later.

In summary, we were able to successfully identify a PKS gene cluster, novel to *Priestia filamentosa,* by assembling its genome, identifying secondary metabolites and screening for the fluorescence of the metabolite produced. The rem gene cluster is known to exhibit several unusual features of the type II PKS involved, a putative MCAT with highest homology to AT domains from modular PKSs. Overall, it was inferred that JURBA-X exhibited fluorescence and antibacterial activity, as a function of these compounds.

## Conclusion

Drought stress affects the growth and development of plants suggesting an utmost need for research identifying new microbes which can help the plant in sustaining such stress. The present study involves the isolation of bacterial isolates from different drought-hit rhizospheres and screening for drought-tolerance using genome sequencing and comparative genomics as a tool. Among the potent ones, *Priestia filamentosa* JURBA-X was identified as an orange-fluorescent *Bacillus* PGPR showing filamentous cellular morphology and mild pink color pigmentation. JURBA-X draft genome was assembled into a 5,113,908 bp genome with 55 contigs and 5,352 protein-coding genes using the Illumina sequencing platform. Due to the presence of genes involved in plant growth promotion traits such as production of siderophore, phosphate solubilization, metabolism of nitrogen and sulfur, and antibacterial (lantibiotics) compound synthesis and osmotic regulation, *P. filamentosa* JURBA-X might be a potent drought tolerant PGPR. The presence of lantibiotics and polyketide genes confers the putative antibacterial activity of JURBA-X. Overall, the consortium of JURBA-X along with other PGPR positive for the antifungal activity could be a potent biocontrol agent against disease-causing phytopathogens and drought stress.

## Ethics approval and consent to participate

Not applicable

## Availability of data and materials

Authors have used open-source tools in this whole analysis. All tool versions have been provided in the methodology. This Whole Genome Shotgun project has been deposited at DDBJ/ENA/GenBank under the accession JAGKQT000000000. The version described in this paper is version JAGKQT010000000.

## Authors’ contributions

KBS generated the idea. MG and KBS cultured the bacteria and did the experimental work. SM and GS performed the computational analysis. SM wrote the first draft of the manuscript. All authors edited and finalized the manuscript.

## Acknowledgments

The authors are thankful to their respective Universities/ Institutes for providing essential facilities and environment for research.

## Supplementary Figure Legends

**Figure-S1:** Phylogenetic characterization of *P. filamentosa* JURBA-X using 16S rRNA nucleotide sequences showcasing the limited resolution of close relatives as witnessed by sporadic distribution of same species.

**Figure-S2:** (A) Phylogenetic analysis based on RemI (B) RemF and (C) RemL proteins, suggesting putative closest relatives of rem cluster

## Supplementary Table Legends

**Table-S1:** Genome-to-genome distance calculation (GGDC) between *P. filamentosa* JURBA-X and its close relative organisms.

**Table-S2:** Comparative ANI calculations between *P. filamentosa* JURBA-X and its close relative organisms.

**Table-S3:** List of annotated genes involved in osmotic stress tolerance, siderophore biosynthesis, auxin biosynthesis, and phosphate-sulfate-nitrogen metabolism in *P. filamentosa* JURBA-X genome.

**Table-S4:** Bacteriocins identified using BAGEL4 in *P. filamentosa* JURBA-X and other related strains

**Table-S5:** Major secondary Metabolites identified using ANTISMASH in *P. filamentosa JURBA-X* and other related strains

