## Supplementary figures and images for "Whole genome sequencing and comparative genomic studies of *Priestia filamentosa* JURBA-X for its drought-tolerance, plant-growth promotion, and fluorescent characteristics"

### Supplementary Figures

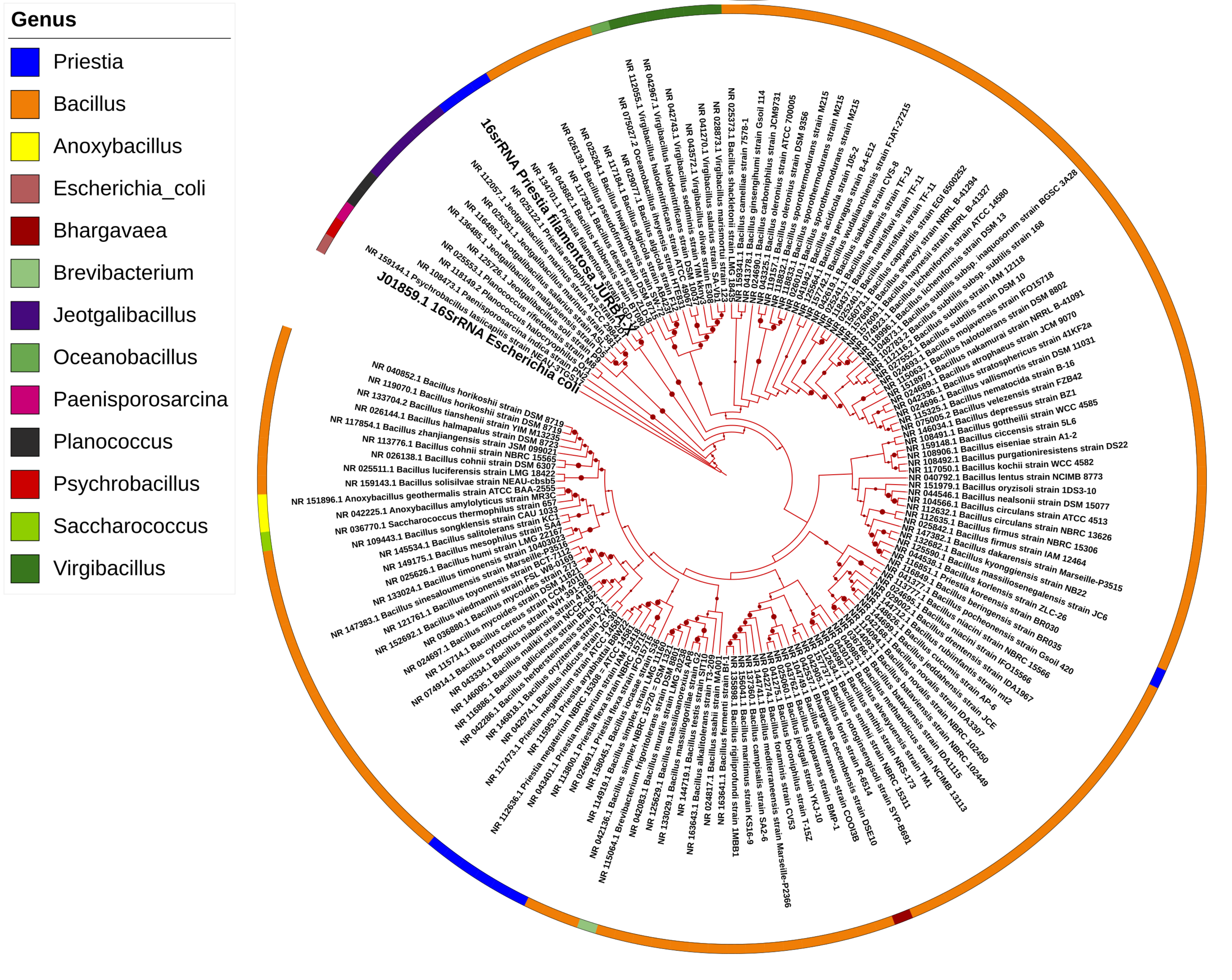

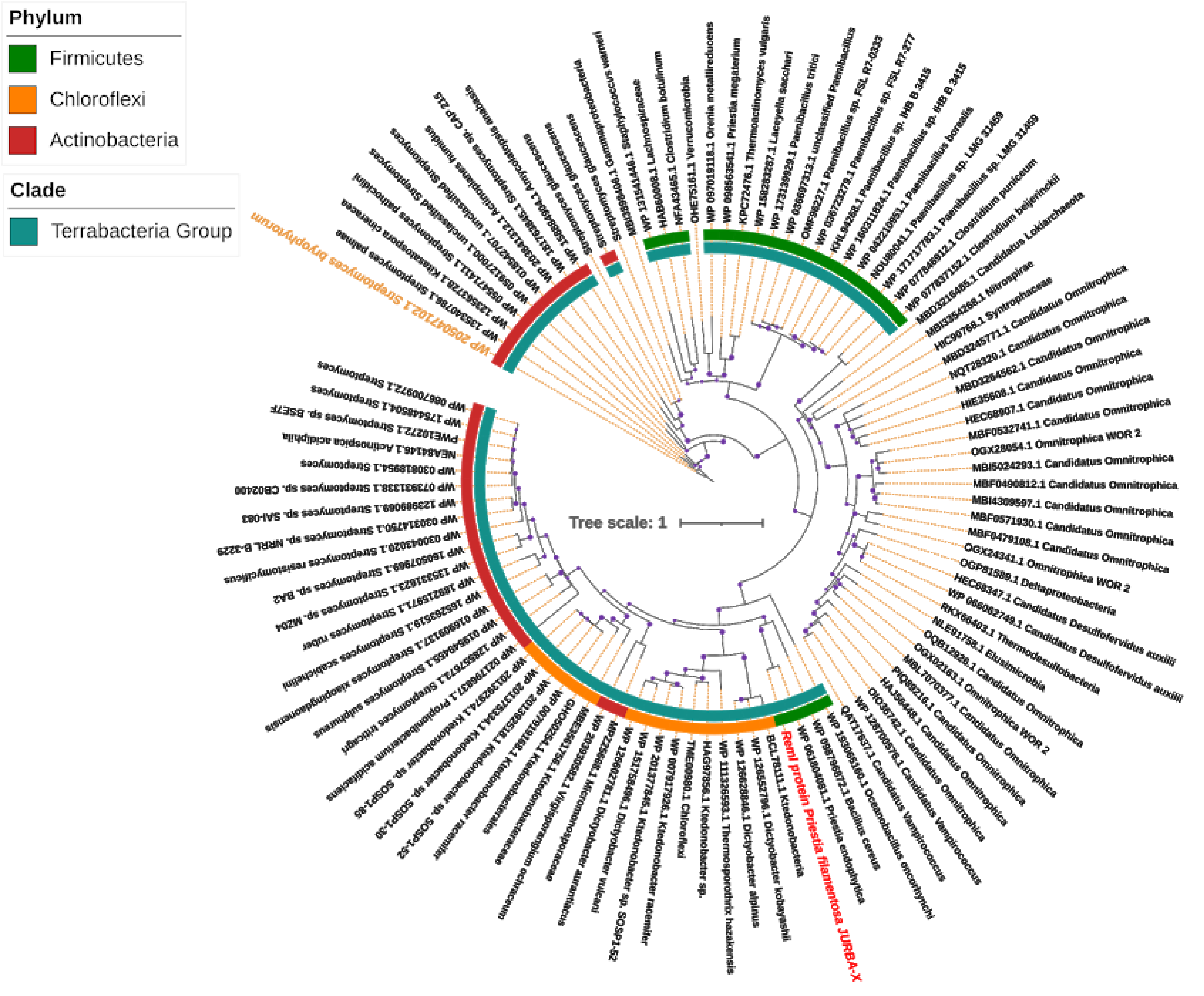

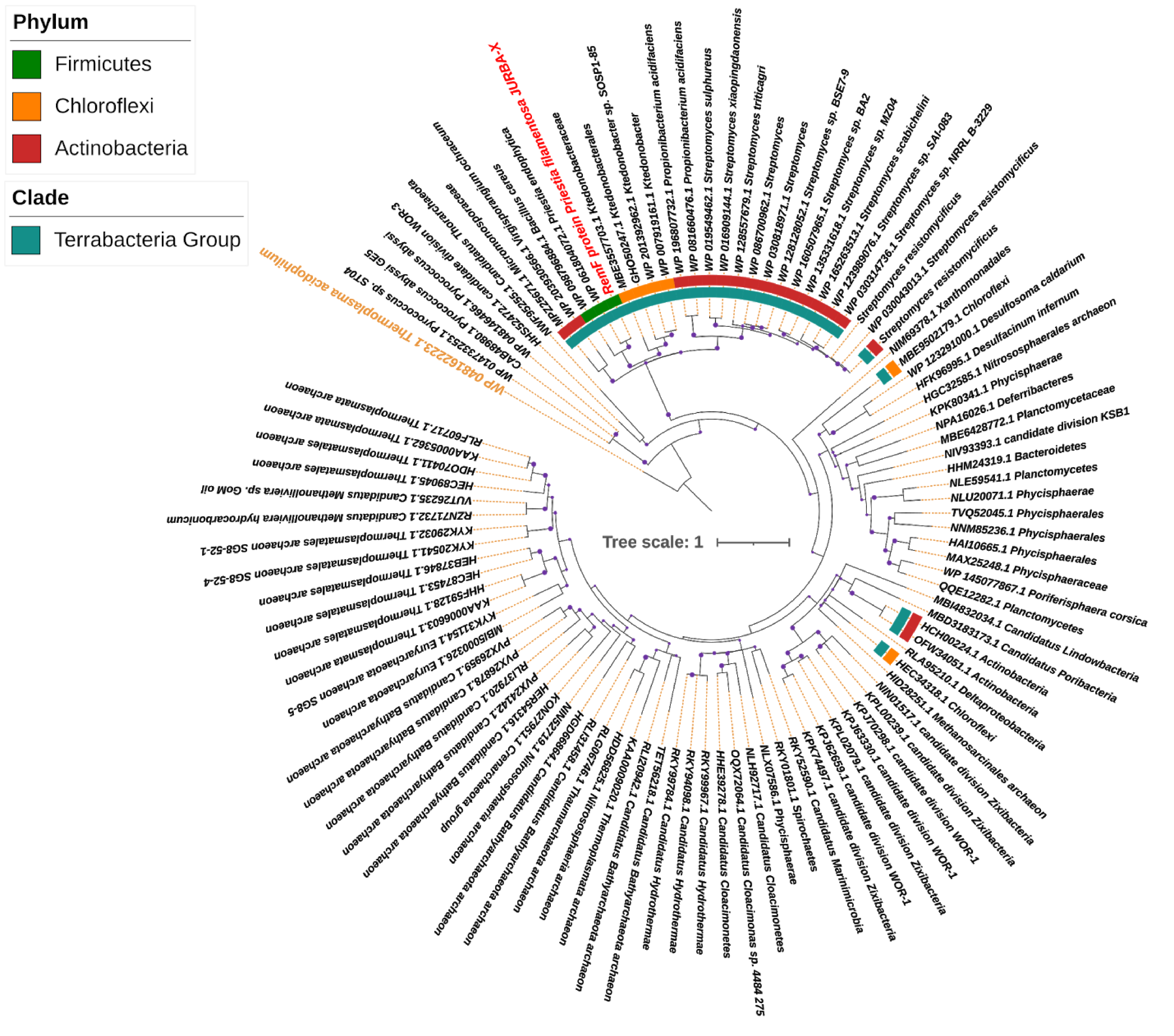

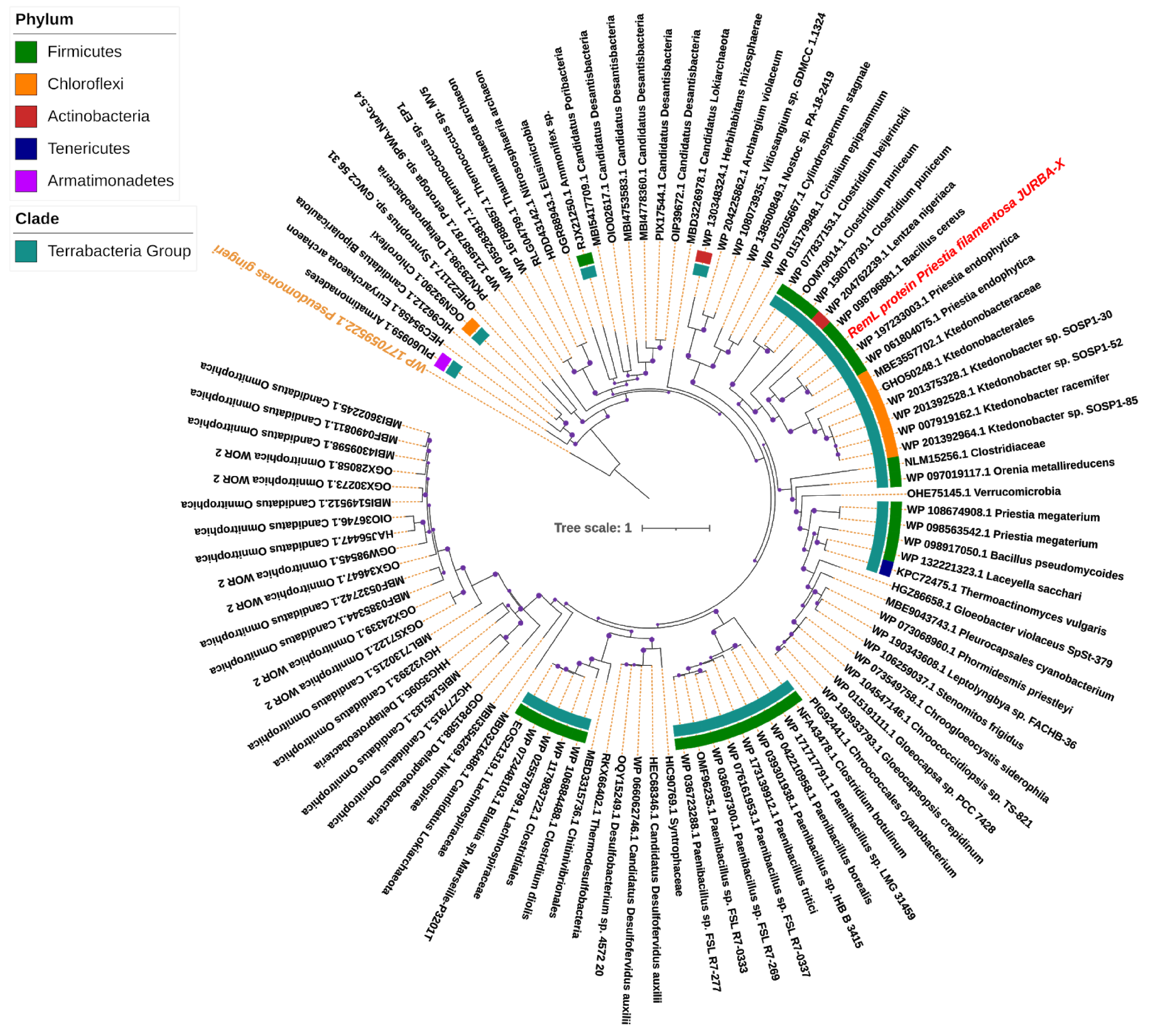
